# Longitudinal single-cell transcriptomics reveals distinct patterns of recurrence in acute myeloid leukemia

**DOI:** 10.1101/2022.01.04.474929

**Authors:** Yanan Zhai, Prashant Singh, Anna Dolnik, Peter Brazda, Nader Atlasy, Nunzio del Gaudio, Konstanze Döhner, Hartmut Döhner, Saverio Minucci, Joost Martens, Lucia Altucci, Wout Megchelenbrink, Lars Bullinger, Hendrik G. Stunnenberg

**Affiliations:** Department of Precision Medicine, University of Campania “Luigi Vanvitelli”, Vico L. De Crecchio 7, 80138 Naples, Italy; Prinses Maxima Centrum, Heidelberglaan 25, 3584 CS Utrecht, The Netherlands; Department of Molecular Biology, Faculty of Science, Radboud University, Radboud Institute for Molecular Life Sciences, Nijmegen, the Netherlands; Charité – Universitätsmedizin Berlin, corporate member of Freie Universität Berlin, Humboldt-Universität zu Berlin, and Berlin Institute of Health, Medical Department, Division of Hematology, Oncology, and Cancer Immunology, Berlin, Germany; German Cancer Consortium (DKTK) and German Cancer Research Center (DKFZ), Heidelberg, Germany; Department of Internal Medicine III, University Hospital of Ulm, Ulm, Germany; Department of Experimental Oncology, European Institute of Oncology, Milan, Italy; BIOGEM, Institute of Molecular Biology and Genetics, Ariano Irpino (AV), Italy

**Author notes:** Corresponding author: Hendrik G. Stunnenberg, Department of Molecular Biology, Faculty of Science, Radboud University, Radboud Institute for Molecular Life Sciences, Nijmegen, the Netherlands, Geert Grooteplein Zuid 28, 6525 GA Nijmegen, The Netherlands. Senior authors (Lucia Altucci, Wout Megchelenbrink, Lars Bullinger, Hendrik G. Stunnenberg). **Competing interests** The authors declare no competing interests.

## Abstract

The heterogeneity and evolution of AML blasts can render therapeutic interventions ineffective in a yet poorly understood patient-specific manner. To gain insight into the clonal heterogeneity of diagnosis (Dx) and relapse (Re) pairs, we employed whole-exome sequencing and single-cell RNA-seq to longitudinally profile two t(8;21) (*AML1-ETO* = *RUNX1-RUNX1T1*), and four *FLT3*-ITD AML cases.

The single cell RNA data underpinned the tumor heterogeneity amongst patient blasts. The Dx-Re transcriptomes of high risk *FLT3-*ITD pairs formed a continuum from extensively changed in the absence of significantly mutational changes in AML-associated genes to rather similar Dx-Re pair of an intermediate risk *FLT3-*ITD. In one high risk *FLT3*-ITD pair, a pathway switched from an AP-1 regulated network in Dx to mTOR signaling in Re. The distinct *AML1-ETO* pairs comprise clusters that share genes related to hematopoietic stem cell maintenance and cell migration suggesting that the Re leukemic stem cell-like (LSC-like) cells probably evolved from the Dx LSC-like cells.

In summary, our study revealed a continuum from drastic transcriptional changes to extensive similarities between respective Dx-Re pairs that are poorly explained by the well-established model of clonal evolution. Our results suggest alternative and currently unappreciated and unexplored mechanisms leading to therapeutic resistance and AML recurrence.

## Introduction

Acute myeloid leukemia (AML) is a malignancy of hematopoietic stem cells or early progenitors resulting from the accumulation of genetic aberrations that disturb key biological processes. Mutations may occur in myeloid progenitor populations, which confer self-renewal capacity to the progenitors^1^. In the past decades, numerous AML associated gene alterations have been identified that can be broadly grouped into four classes^2^. Class I comprises the mutations that activate signal transduction pathways and induce the proliferation or survival of HSPCs, such as *FLT3*^3,4^, *NRAS*/*KRAS*^5^ and *KIT*^6^. Class II consists of mutations or fusions in genes coding for transcription factors that are required for hematopoietic maturation, like *AML1-ETO* (*RUNX1-RUNX1T1*)^7^ and *CEBPA*^8^. Class II aberrations happen during early hematopoiesis and initiate leukemia, while Class I aberrations take place in later stages and cause leukemia expansion. Class III consists of epigenetic regulators like *IDH1/2, TET2, DNMT3A* and *ASXL1*, whereas class IV consists of tumor suppressor genes, such as *TP53*.

Despite that current chemotherapies efficiently induce complete remission, AML patients frequently suffer from relapse and have low overall 5-years survival rates^9–11^. Recurrence can emerge as a result of the expansion of pre-existing chemo-resistant subpopulations or by acquiring novel chemo-resistant subpopulations due to genomic altereations^12^. The advent of single-cell RNA sequencing provides revolutionary opportunities to assess the heterogeneity of cancer populations at the single-cell level and explore the transcriptional features of individual cell types, such as subpopulations contributing to the relapse. However, few longitudinal studies^13,14^ focused on analyzing pair-wise samples from AML patients, at first diagnosis and relapse.

Here, we applied single-cell RNA sequencing to analyze dynamic changes of gene expression between AML samples at diagnosis and at relapse. We profiled 5 612 high-quality cells at diagnosis and relapse from 6 AML patients, n=2 low risk cases with t(8;21) (*AML1-ETO*), n=1 intermediate and n=3 high risk AML cases with *FLT3-*ITD. Whole-exome sequencing (WES) was used to study the acquired genomic mutational profile. Our single cell RNA study uncovered extensive inter- and intra-heterogeneity amongst *AML1-ETO* and *FLT3-*ITD pairs at diagnosis (Dx) and relapse (Re). Our study provides novel insights into recurrence and unveisl vulnerabilities that could serve as new entry points for targeting relapse AMLs.

## Methods

### AML samples and cell preparation

We processed 6 paired Dx-Re bone marrow aspirates from adult AML patients, with *AML1-ETO* (n=2 low risk cases) or *FLT3-*ITD (n=1 intermediate and n=3 high risk). Patients characteristics are summarized in Supplemental Table 1. CD33/CD34+ cells were sorted into 384-well plates and stored at -80°C.

### Single cell SORT-seq

SORT-seq^15^ is based on the integration of single cell FACS sorting (Fluorescence-Activated Cell Sorter) with the CEL-Seq2 protocol^16^. Single cell libraries were paired-end sequenced on an Illumina NextSeq500 at an average depth of ∼30M reads per library.

### Fusion genes detection

To quantify the reads per gene and detect fusion genes from bulk RNA-Seq, sequence libraries were aligned to Gencode v37 reference genome version hg38 using STAR-Fusion v1.10.0^17^ in 2-pass mode, with parameters -*-CPU 12 --FusionInspector validate --examine_coding_effect --denovo_reconstruct*.

### Whole-exome sequencing

WES libraries were generated as previously described^18^. Diagnosis and relapse samples were compared with samples collected at CR (Complete Remission).

### Pseudo-time trajectory analysis

We used Monocle3^19,20^ for pseudo-time analysis with default parameters, to assess the trajectories within the pairs. We used the DEGs obtained from Seurat 3.0^19^ to plot the dynamic changes of gene expression along the trajectories.

### Definition of leukemic stem cells and cycling genes

The 17-gene leukemic stem cell (LSC17) score was calculated based on the equation by Ng et al.^21^. Cell cycle phase scores were calculated using Seurat 3.0 function *CellCycleScoring* with default parameters.

## Results

### Whole exome- and gene fusion analysis points to limited clonal rearrangements between Dx and Re

Clonal expansion and evolution is a major determinant of AML relapse^22^. To identify the genomic landscape at Dx and Re, we performed whole exome sequencing analysis (WES) and gene fusion detection based on bulk RNA-sequencing. We detected 4 up to 26 somatic mutations in the Dx and Re pairs (Figure 1A, Supplemental Table 2). This analysis confirmed the presence of an inframe insertion in the juxtamembrane domain (JMD) between amino acid 583 and 611 in all four patients diagnosed with *FLT3-*ITD as well as *AML1-ETO* fusion transcripts in the *AML1-ETO* patients (Figure 1A-B, Supplemental Table 2). Other AML-associated somatic variants, such as *NPM1, WT1, CEBPA, IDH1, NRAS* and *DNMT3A* were detected for the *FLT3*-ITD patients, often in a patient-specifc manner. For both *AML1-ETO* patients, the WES analysis revealed a *KIT* mutation that is associated with poorer prognosis and increased risk of relapse^23,24,25^.

**Figure 1.**
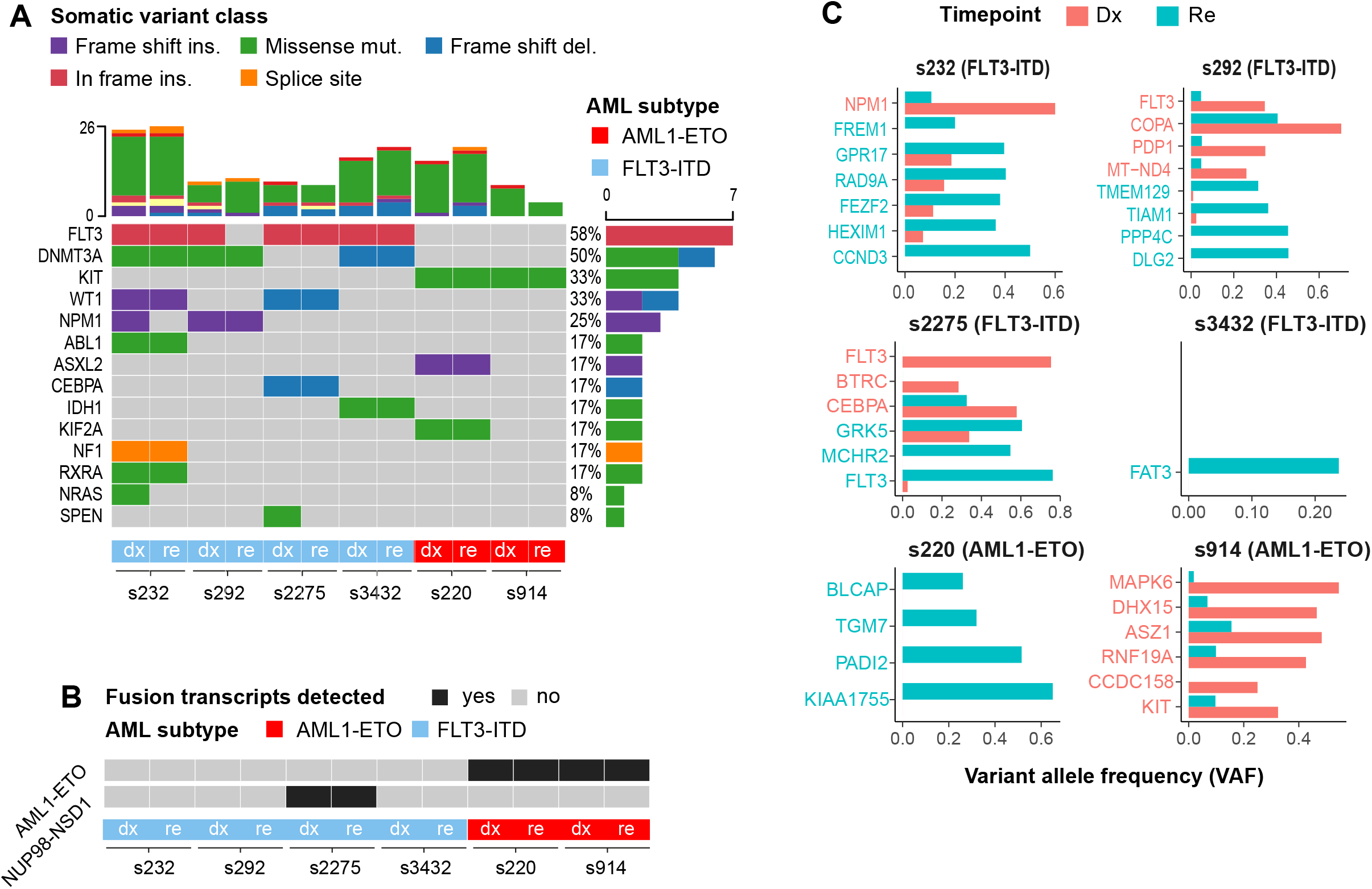
Whole exome- and gene fusion analysis between Dx and Re. (A) Oncoplot from WES showing 14 selected somatic mutations across 6 patients (red: n=2 AML1-ETO; blue: n=4 FLT3-ITD). Mutations with at least 5 reads on ALT allele and VAF ≥ 0.05 are presented. Vertical bars depict the number of mutations detected per sample; horizontal bars depict the (relative) frequency of a particular mutation. (B) Gene fusions detected from bulk RNA-seq. (C) Mutations with a VAF ≥ 0.2 at Dx or Re for which the VAF changed significantly. For all bars, p < 0.05, Fisher’s exact test with Benjamini-Hochberg correction. Red: mutations more abundant at Dx. Blue: mutations more abundant at Re.

Next, to identify clonal rearrangements that may have led to disease relapse, we screened for somatic mutations with a significantly altered variant allele frequency (VAF) between Dx and Re (VAF ≥ 0.2 and p < 0.05, Fisher’s exact test; methods). For patient s232, WES and PCR analysis revealed two distinct *FLT-*ITD mutations in Dx sample, one of which one was lost at Re (p=1.0 × 10^−3^; Fisher’s exact test; Figure 1A, Supplemental Table 2). WES analysis further revealed the presence of 4bp insertion in *NPM1* (mutation type A^26^) at Dx, that was decreased at Re (p=8.2 × 10^−3^) as well as a lowly abundant missense mutation in the *NRAS* gene at Dx (VAF=0.087) that was not detected at Re (VAF=0; p=2.0 × 10^−4^; Figure 1C). For patient s2275, the WES data showed considerably shorter tandem duplications at relapse compared to diagnosis (p = 4.6 × 10^−41^), which were confirmed by PCR (Supplemental Table 3) as well as the presence of *NUP98-NSD1* fusion transcripts at Dx and Re. We further detected a 4bp insertion in *NPM1* and a missense mutation in DNMT3A that are retained between Dx and Re in patient s292. For patient s3432, WES and PCR showed a retention of the *FLT3-*ITD, both in the insertion location and allelic ratio. Somatic mutations in *FAT3* (VAF= 0.238, p = 4.3×10^−8^), *ITGB7* (VAF= 0.165, p = 1.32 × 10^−6^), *UBA2* (VAF = 0.117, p=6.32 × 10^−3^) and *SLC4A3* (VAF = 0.135, p= 6.6 ×10^−3^) were significantly gained in the Re sample (Figure 1C and Supplemental Table 2). Two distinct *KIT* mutations (VAF = 0.325; VAF = 0.138, respectively) were detected in patient s914 at Dx, both of which were significantly reduced at Re (p < 4.7 × 10^−7^). Finally, other somatic mutations that have not been implicated with AML in the Catalogue of Somatic Mutations in Cancer^27^ (COSMIC), were lost or gained in all pairs (Figure 1C; methods).

To summarize, we confirmed the presence of *FLT3-*ITDs and *AML1-ETO* in four and two patients respectively. Additional somatic aberrations in AML-associated genes were patient-specific. *FLT3-*ITD mutations were altered in two patients and in one patient, one of the two *FLT3-*ITD mutations was lost at Re. For patient s232, a *NPM1* mutation was detected at Dx, but lost at Re. Finally, we observed a significant reduction in two distinct *KIT* mutations in patient s914 between Dx and Re.

### Single cell transcriptomics reveals distinct AML-phenotypes at Dx and Re

Next, to better understand the transcriptional phenotypes, their differences and possible mechanisms that led to disease progression, we profiled bone marrow cells obtained at Dx and after Re using single cell transcriptomics. In brief, single CD33^+^ or CD34^+^ bone marrow cells were FACS-sorted into 384-well plates following the SORT-seq method^15^ we acquired 5 612 single cell profiles, in which 4 129 unique transcripts from 1 678 genes were detected on average (Supplemental Figure 1A, methods).

After normalization, cells were clustered and visualized using the uniform manifold approximation and projection^28^ (UMAP). *AML1-ETO* vs *FLT3-*ITD samples are separated by UMAP1 and Dx-Re pairs cluster relatively close together (Figure 2A-B). Nevertheless, considerable heterogeneity between and within pairs exists (Figure 2B). Strikingly, Dx-Re cells of *FLT3*-ITD patient s232 cluster in close proximity suggesting minor phenotypic and molecular alterations, eventhough this patient lost *NPM1* and *NRAS* mutation at Re. In contrast, Dx cells of patient s3432 are completely separated from Re cells, athough one mutation in the *FAT3* gene was detected in Re (VAF=0.238) (Supplemental Table 2). Similarly, the Dx and Re cells of *AML1-ETO* patient s220 constitute distinct clusters, but only gained mutations in genes that are not associated with AML (Figure 1C). Unexpectedly, patient s914 had a significant loss of two *KIT* mutations between Dx (VAF=0.325 and 0.138) and Re (VAF= 0.097 and 0) that resulted in relatively small transcriptional alterations.

**Figure 2.**
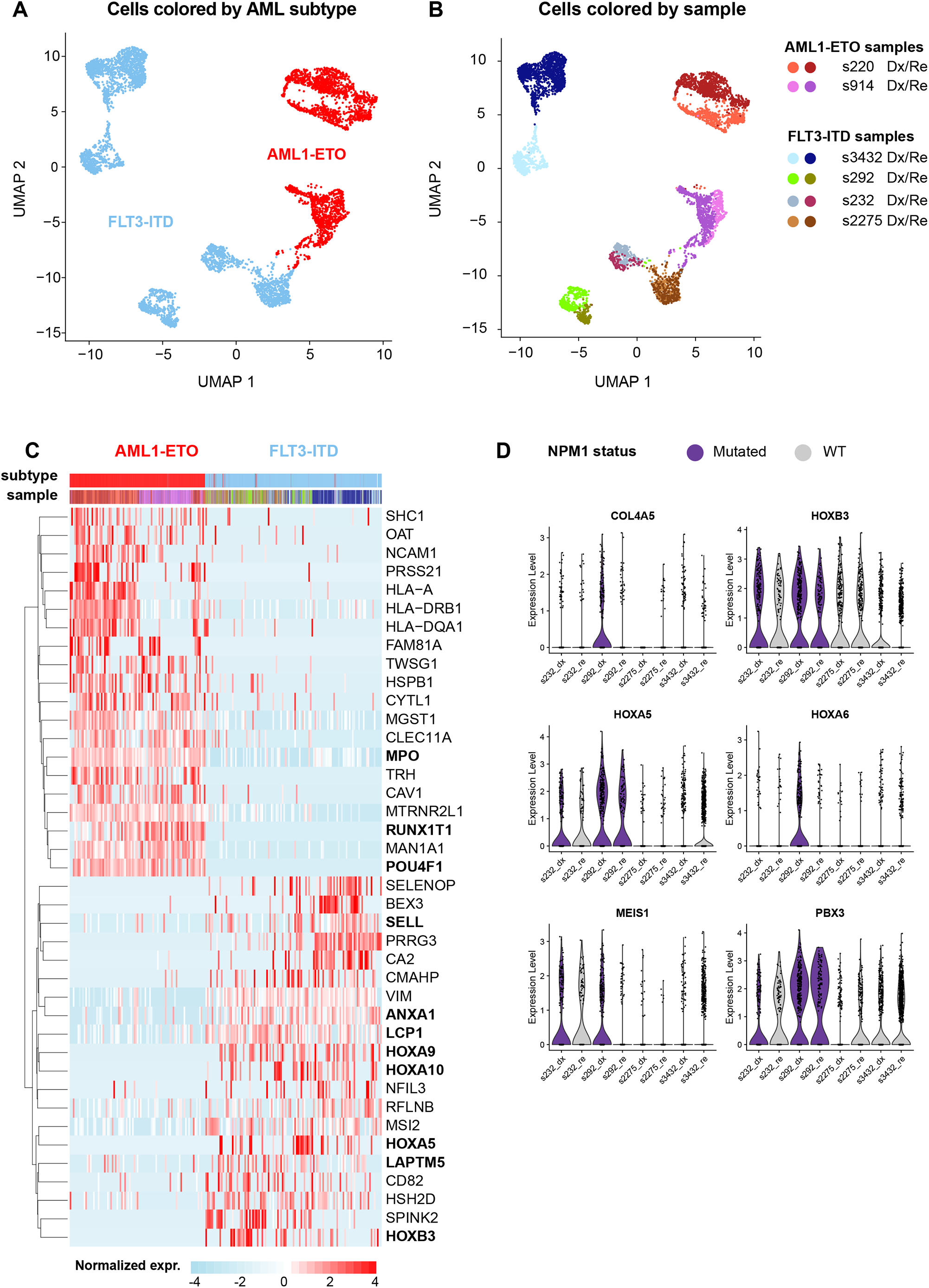
Single cell transcriptomics reveals distinct AML-phenotypes. (A) UMAP of the six AML pairs, colored by primary mutation (red: *AML1-ETO*; blue: *FLT3*-ITD); (B) UMAP colored by sample; (C) Heatmap showing the top 20 marker genes per primary mutation **D**, Violin plots depicting gene expression at known NPM1 target genes in *FLT3*-ITD samples with- and without *NPM1* mutation.

To further verify the quality of our single cell data, we looked for gene signatures that discriminated *AML1-ETO* or *FLT3-*ITD patients. These signatures include well-established *AML1-ETO* markers, like upregulation of the transcriptional co-repressor *RUNX1T1* (aka *ETO*), the transcription factor *POU4F1*^29^ and the myeloid differentiation protein *MPO*^30^ (Figure 2C, top). *FLT3*-ITD samples on the other hand are characterized by *VIM, ANXA1, MSI2, LAPTM5*. Other genes tend to be overexpressed only in a subset of the samples: *HLA* genes are overexpressed in *AML1-ETO* patient s220, but not in s914. In the *FLT3-*ITD samples, *HOXA5* and *HOXB3* genes that are overexpressed in *NPM1*-mutated AML^31^, appear overexpressed in a patient-specific manner (Figure 2C, bottom). Closer inspection of these and other *NPM1*-marker genes showed that these genes are indeed signicantly higher expressed in the FLT3-ITD samples with an additional *NPM1* mutation (*NPM1*^mut^) compared to *NPM1*^WT^ samples (FC > 1.5 and p < 6.0 × 10^−15^; Figure 2D). Notably, *HOX*-genes are also highly expressed in *FLT3-*ITD patient s2275. In these samples, we detected a *NUP98-NSD1* fusion gene that is characterized by upregulation of *HOXA* and *HOXB* genes^32^ (Figure 2D).

In summary, single cell transcriptomics showed distinct clustering of *AML1-ETO* vs *FLT3-*ITD patients. Differential analysis confirmed upregulation of well-established marker genes as well as elevated expression of *HOX* genes in *NPM1*^mut^ and the *NUP98-NSD1* positive *FLT3-*ITD samples. On a global level, the transcriptional changes between Dx and Re are poorly explained by mutations in coding regions of AML-associated genes. To gain a deeper understanding of the mechanisms underlying these changes, we subsequently performed an indepth analysis of Dx-Re pairs per AML-subtype and in a patient-specific setting.

### Dx-Re transcriptomic changes are patient specific

Given this high intra- and inter-patient heterogeneity, we focused on the Dx-Re differences per patient in the remainder of this study. For this, we separated the UMAPs of the *FLT3*-ITD and *AML1-ETO* patients (Figure 3A-B) and computed the differentially expressed genes between the Dx-Re pairs per patient. This analysis reinforced the notion that the differences in transcription between Dx and Re are highly patient specific (Supplemental Figure 1B-C, Figure 3C-D).

**Figure 3.**
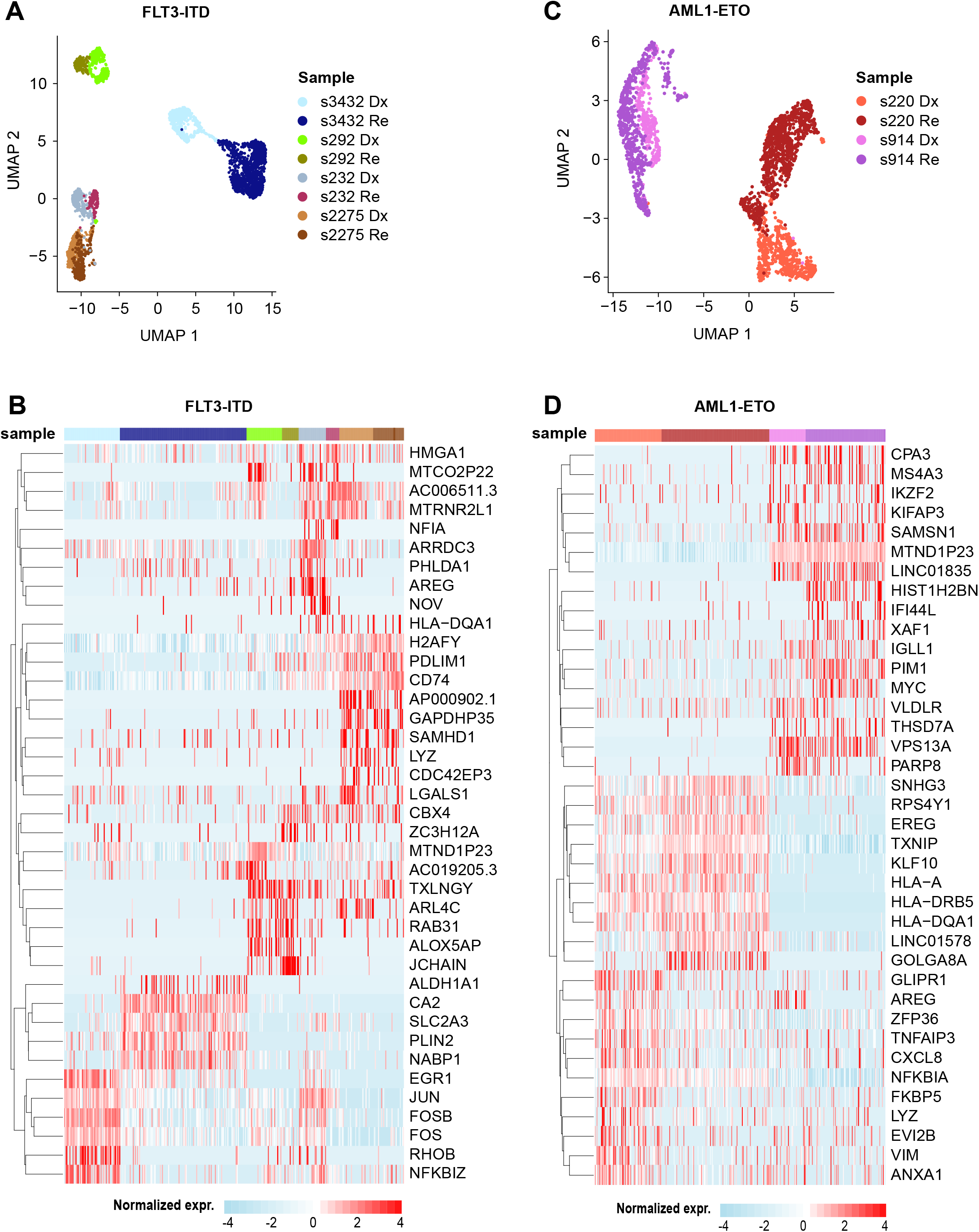
Single cell transcriptomics reveals heterogeneity amongst patients. (A) UMAP of the four sample pairs with a *FLT3*-ITD, colored by sample (red: Dx; blue: Re); (B) Heatmap displaying the top 5 marker genes per sample (*FLT3*-ITD); (C) UMAP of the two *AML1-ETO* sample pairs, colored by sample; (D) Heatmap displaying the top 10 marker genes per sample.

The *FLT3*-ITD patients show a modest separation between the Dx and Re samples of patient s232 (Figure 3A). Cluster analysis revealed two clusters at diagnosis (cluster 1-2) and one at relapse (cluster 3, Supplemental Figure 2A). Re cells lost expression of members of the *AP-1* transcription factor, like *FOS, FOSB* and *ATF3* that were highely expressed in Dx cluster 1 (Supplemental Figure 2B). Gene ontology (GO) analysis confirmed significant loss of expression for these and other genes involved in *AP-1*/*ATF-2* related transcription at Re (Supplemental Figure 2C). Furthermore, we evaluated the expression level of genes involved in PI3K/AKT/mTORC pathway, in which mTORC1 controls ribosomal biogenesis and protein translation^33^. We found the targets of mTORC1, like *RPS6KB1* and *EIF4E*, were differentially expressed in Re (Supplemental Figure 2D), suggesting a pathway shift from AP-1 to mTORC1. Besides, we observed the upregulation of the upstream *K/NRAS* genes in Re, which may be markers for diagnosis/ prognosis and treatment target.

The UMAP for patient s292 showed 3 distinct clusters (Supplemental Figure 3A). DEG between Dx clusters 1 and 2 revealed *IDH1*, an enzyme in the TCA cycle, and *RAB31* involved in membrane fusion and exocytosis in clusters 1, whereas MPO and PROM1, markers for GMP cells, are differentially expressed in cluster2 (Supplemental Figure 3B-C). Cells in cluster 3 originate from the Re sample and overexpressed genes like *DDIT4*^34^, *PIM3*^35^ and *CD74*^36^ were previously associated with poor prognosis (Supplemental Figure 3B). GO analysis indicated regulation of cell death and apoptotic process terms in cluster 3 (Supplemental Figure 3C). For patient s2275, the single cell expression analysis detected 5 clusters. Cluster 1 mainly originated from Dx cells, whereas cluster 5 almost entirely consisted of Re cells. Clusters 2-4 however were a mixture between Dx and Re cells (Supplemental Figure 4A-B). DEGs revealed few differences between cluster 1 and 5, such as *RNU4ATAC* and *RYBP* involved in RNA biosynthesis and metabolics that are differential expressed in cluster 1 (Dx) (Supplemental Figure 4B-C), whereas *ITM2A* and *CLEC12A* for leukocyte activation and *LDHA* for ribonucleotide metabolics are differentially expressed in cluster 5 (Re) (Supplemental Figure 4B-C). The minor differences between Dx and Re is consistent with the fact that AML-associated mutations, such as *WT1, CEBPA* and *NUP98-NSD1* are retained at Re (Figure 2C, Supplemental Table 2).

**Figure 4.**
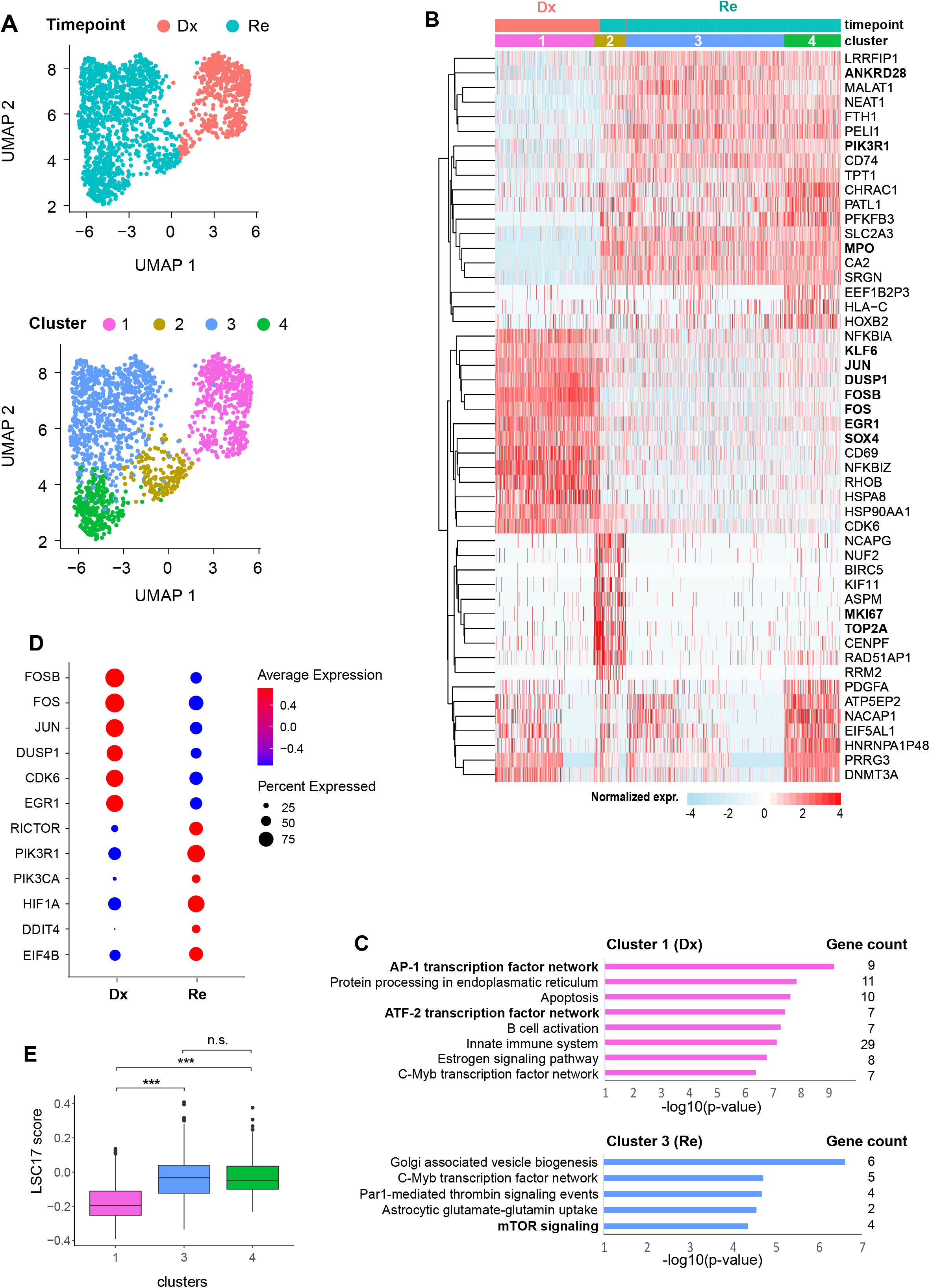
Pathway switch between AP-1 and RAS signaling in high risk *FLT3*-ITD (s3432) (A) UMAP of Dx and Re cells for *FLT3*-ITD patient s3432 colored by timepoint (top) or cell cluster (bottom). (B) Heatmap displaying the top 10 cluster marker genes. Color represents row normalized expression values. (C) Overrepresented GO terms (category: biological pathway) in cluster 1 (Dx) and 3 (Re). P-values: hypergeometric test (BH-corrected). (D) The expression of genes related to AP-1 transcription factor network and RAS signaling pathway in each timepoint. (E) Calculation of LSC17 score for each cluster, and p-value was calculated using Student’s t-test. * p < 0.05, ** p < 0.01, *** p < 0.001.

The Dx and Re cells of patient s3432 formed distinct clusters that are highly separated from each other and the other *FLT3*-ITD patients (Figure 3A). Cluster analysis detected four groups of cells that largely separated Dx (cluster 1) from Re cells (cluster 2-4; Figure 4A). Cluster 1 had a characteristic gene signature of transcription factors involved in proliferation and cell growth (e.g., *JUN, FOS, FOSB, EGR1, SOX4* and *KLF6*) that were significantly downregulated in the relapse clusters (Figure 4B-C). The Re-specifc clusters 3-4 upregulated genes involved in the RAS/mTORC pathway, such as *ANKRD28* and *PIK3R1*, whereas cluster 2 is hallmarked by cell cycle related genes, such as *TOP2A* and *MKI67*. Pathway enrichment analysis confirmed the overrepresentation of AP-1/ATF2 transcription factors in cluster 1 (Dx) and additionally revealed upregulation of genes involved in mTOR signaling, like *RICTOR, PIK3R1* and *HIF1A* in cluster 3 (Re; Figure 4C-D). This suggests a pathway switch from AP-1 in the diagnosis cells towards mTOR in the relapse cells. We further observed that *KRAS* and *NRAS*, genes upstream of mTORC, were also overexpressed in the Re sample (Supplemental Figure 5A). Interestingly, cluster 4 in relapse is characterized by elevated exocytosis (Supplemental Figure 5B) and increased expression of genes related to Tim-3-galectin-9 Secretory Pathway (e.g. *ADGRL1, HAVCR2* and *LGALS9*) that protect AML cells against from the host immune system in an mTOR dependent manner^37^ (Supplemental Figure 5C), in particular from NK- and T-cell action. Finally, the leukemia stem cell (LSC) score, a 17-gene signature (LSC17) that correlates with aggressiveness of the leukemia and a poor outcome^21^ was significantly higher in the Re clusters 3 and 4 compared to the Dx cluster 1 (Figure 4E).

### Leukemic Stem Cell-like cells in *AML1-ETO*

In line with elevated expression of the *RUNX1T1* (aka *ETO*) and the well-known target gene *POU4F1* (Figure 3C), *AML1-ETO* fusion transcripts were detected in the Dx and Re samples of both patients (Figure 1A, Supplemental Table 2). WES analysis had further revealed that both patients suffered from one (s220) or two KIT mutations at time of Dx that were retained for patient s220 at Re, but largely or exclusively lost for patient s914 (Figure 1C, Supplemental Table 2). Surprisingly, UMAP and DEG analysis revealed signficantly larger transcriptional changes for patient s220 compared to s914 (Figure 3C, Supplemental Figure 1C). Possibly, these transcriptional changes are induced by the somatic mutations in genes that are not widely associated with AML, like *BLCAP, TGM7, PADI2* and *KIAA1755* (Figure 1C). Higher *MPO*, a marker for granulocyte/monocyte progenitors (GMPs) expression^30^ within both the *AML1-ETO* patients (Figure 3C) implies that most cells are arrested at a “GMP-like” stage.

Analysis on Dx-Re showed that the number of DEGs shared between these two *AML1-ETO* patients is minimal as for the *FLT3-*ITDs (Supplemental Figure 1C). Therefore, we performed an in-depth analysis on the transcriptional dynamics between Dx and Re separately for these two patients. Focussing on patient s914 first, the synergic oncogenes (*PIM1* and *MYC*^38^) responsible for tumorigenesis were co-differentially expressed at Re compared to Dx. Cluster analysis revealed five groups of cells (Figure 5A-B) and a small cluster of scattered cells that expressed signatures of progenitors (*CD34*), erythrocytes (*HBB*), monocytes (*LYZ*), B-cells (*MSA41*) and cell cycle related genes (*TOP2A, MKI67*) (Supplemental Figure 6) likely resulting from ambient RNA or cell doublets and hence were discarded in subsequent analyses.

**Figure 5.**
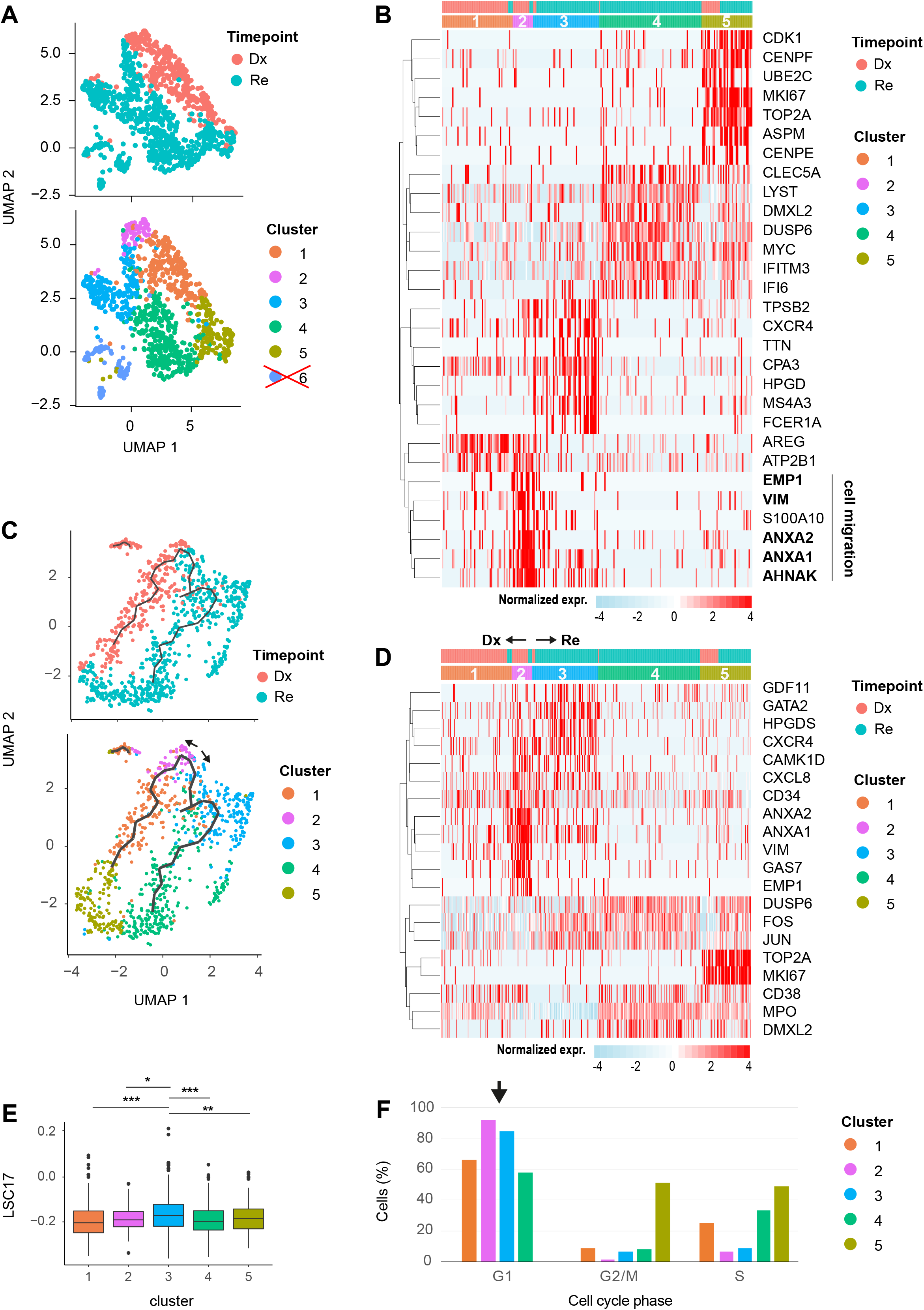
Putative LSCs detected in *AML1-ETO* pair (s914) (A) UMAP of Dx and Re cells for *AML1-ETO* patient s914, colored by timepoint (top) and cell cluster (bottom). Cells in cluster 6 express ambiguous marker genes, and may be doublets or contaminated by ambient RNA and were discarded (see also Supplemental figure 6). (B) Heatmap depicting the top 7 cluster markers. Color represents row normalized expression values. (C) Pseudo-time trajectory colored by timepoint (top) or cell cluster (bottom). (D) Heatmap showing representative genes per cluster. (E) LSC17 scores per cluster. * p < 0.05, ** p < 0.01, *** p < 0.001, Student’s t-test. (F) Barplots depicting the relative cell abundance per cell cycle phase (inferred from marker gene expression) for each cell cluster. Arrow: cells in cluster 2 and 3 predominantly reside in the G1 phase.

Cluster 1 mainly consist of Dx cells and differentially expressed genes for differentiation and resistance to apoptosis, like *AREG*^39^. Interestingly, cells in cluster 2 express *CD34* as well as genes involved in cell migration (*ANXA1*^40^, *ANXA2*^41^, *VIM*^42^ and *EMP1*^43^) but lacked the expression of *MPO* (Figure 5B,D). To investigate whether and from which Dx cluster these potential Re LSCs originate, we aligned cells in pseudo-time based on the gradient of transcriptional differences using Monicle3. This trajectory analysis suggested a continuous transition between the Dx and Re sample (Figure 5C). Cells in cluster 2 and 3 differentially expressed genes for hematopoietic stem cell maintenaince (*GDF11*^44^, *GATA2*^45^) and differentiation (*GAS7*^46^, *CAMK1D*^47^) markers as well as *CD34* (Figure 5B,D), indicating cluster 2 and 3 are the putative starting points of this trajectory. Besides, cluster 2 and 3 overexpressed genes *CXCR4*^48^ and *CXCL8*^49^ for tumor microenviroment (Figure 5B,D). In line with those findings, we calculated the LSC17- and cell cycle scores for all clusters. We observed that cells in Dx cluster 3 has the highest LSC17 score followed by Re cluster 2 (Figure 5E). Moreover, cells from cluster 2 and 3 mainly reside in the G1 phase of the cell cycle (Figure 5F). Interestingly, the trajectory suggest that these cells differentiate into a population of cells that display *DUSP6* and AP-1 related genes like *JUN* and *FOS* in the Re-specific clusters 3 and 4 (Figure 5D).

UMAP shows that s220 cells separate according to Dx and Re which partitioned into 9 clusters (Figure 6A). Clusters 1-4 contained Dx cells that were enriched for *CXCL8* and *CXCR4*, genes associated with the interaction between leukemia blasts and stromal cells^48,49^. Clusters 5-9 exclusively contained Re cells and were marked by expression of *LOXL1* and *FAM81A* (Figure 6B). Cell cycle-related genes (*MCM6, TOP2A, MKI67*) were highly expressed in cluster 1 and 9. Cluster 4 (Dx) and 5 (Re) are in close proximity to each other and share marker genes, such as *CAMK1D, GAS7, ANXA1*/*2, VIM* and *CD34* (Figure 6B, Supplemental Figure 7A) suggesting that they are LSCs.

**Figure 6.**
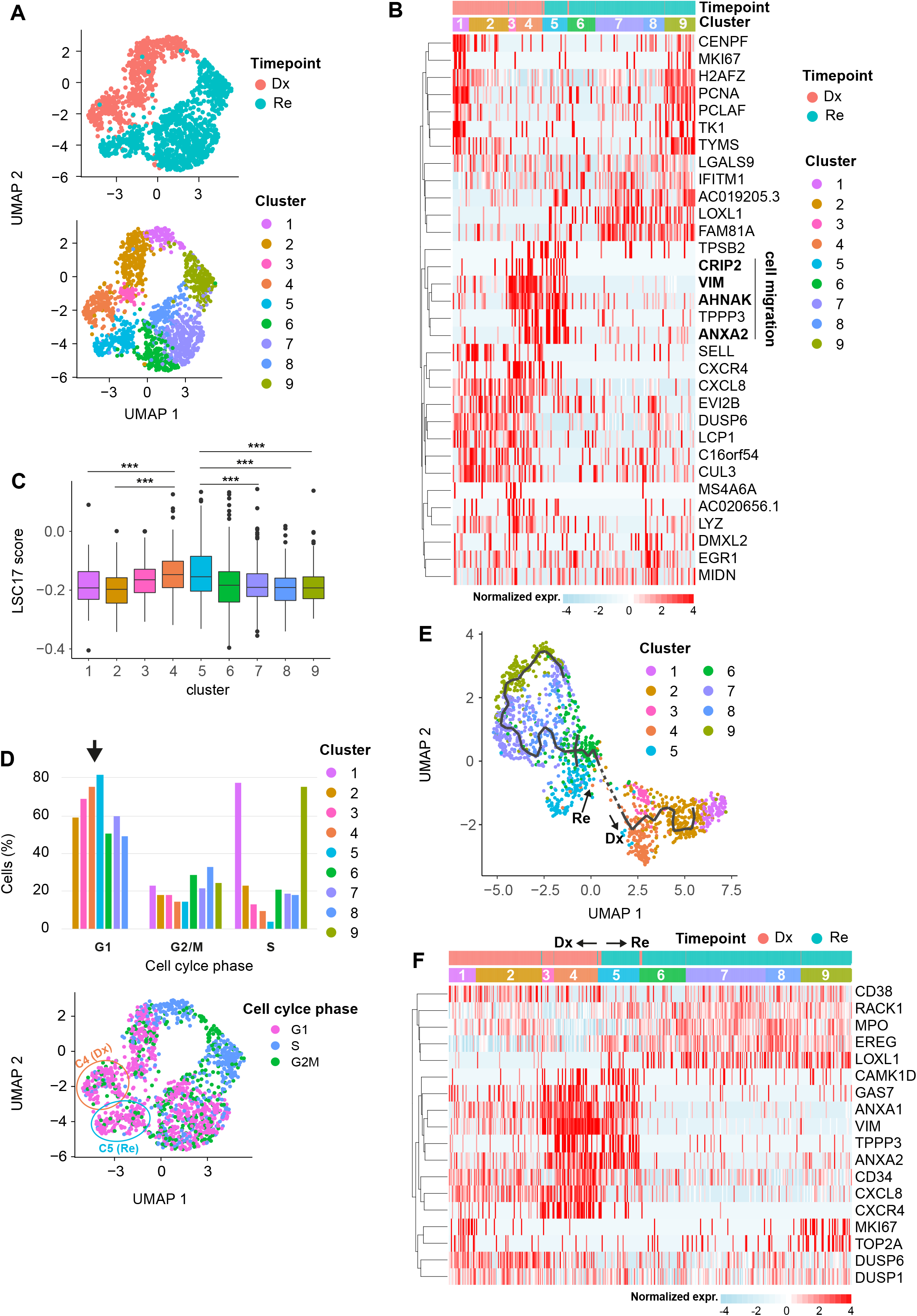
Putative LSCs detected in *AML1-ETO* pair (s220) (A) UMAP of Dx and Re cells for *AML1-ETO* patient s220, colored by timepoint (top) and cell cluster (bottom). (B) Heatmap depicting the top 5 marker genes per cluster. Color represents row normalized expression values. (C) LSC17 scores per cluster. * p < 0.05, ** p < 0.01, *** p < 0.001, Student’s t-test. (D) top: Barplots depicting the relative cell abundance per cell cycle phase (inferred from marker gene expression) for each cell cluster. Arrow: cells in cluster 4 and 5 predominantly reside in the G1 phase. Bottom: UMAP colored by cell cycle phase. (E) Pseudo-time trajectory colored by cell cluster (F) Heatmap depicting representative marker genes per cluster/inferred timepoint.

### Alternative “branching” from Re and Dx LSC-like cells in *AML1-ETO*

Given the high similarities between clusters 4 and 5 and their elevated *CD34* expression, we hypothesized that these clusters might be enriched in LSCs. Analysis showed that these clusters indeed have the highest LSC17-score and contain cells that reside predominantly in the G1 cell cycle phase (Figure 6C-D). To better understand the transcriptional dynamics of cell populations originating from these LSCs, we applied pseudo-time gene expression analysis (Figure 6E). This analysis reveals a trajectory starting from the presumed LSCs cluster 4 and 5 towards more differentiated cells that predominantly reside in the S-phase of the cell cycle and exhibit elevated expression of genes like *TOP2A* and *MKI67* (Figure 6D-F). For the Dx branch, genes involved in self-renewal that impede differentiation (*GAS7* and *CAMK1D*) or are associated with cell migration (*TPPP3, VIM, ANXA1*/*2*) had elevated expression in cluster 4. We hypothesized that all other clusters of cells originate from this presumed Dx LSC population. Indeed, we observed a downregulation of these markers when cells are traced along the trajectory from cluster 4 to cluster 1 which is consistent with their differentiation into more mature myelod cells. Furthermore, *DUSP1* and *DUSP6*, genes required for cell differentiation and proliferation were upregulatd as cells ‘moving away’ from cluster 4 along the Dx branch (Figure 6F). In the Re branch, *TPPP3, VIM, ANXA1*/*2, GAS7* and *CAMK1D* were upregulated in cluster 5 to a similar extent as in cluster 4. Compared to the more gradual downregulation in the Dx branch, these markers were largely lost when cells “branched” from cluster 5 to cluster 6 (Figure 6F). The Re trajectory (cluster 5 towards cluster 9) is hallmarked by upregulation of numerous genes required for differentiation, leukemia progression and chemo-resistance, including *RACK1*^50^, *EREG*^51^ and *LOXL1*^52^ (Figure 6F). Another striking difference between the Dx and Re is that genes associated with the tumor microenvironment, the interaction between stroma cells and leukemic blasts (*CXCR4* and *CXCL8*) were lower expressed in cluster 5 (Re) compared to cluster 4 (Dx, Figure 6B). Gene Ontology analysis further revealed up-regulated genes in Dx enriched with terms associated with immune- and inflammatory response, whereas translation and biosynthesis related processes were highly enriched in Re (Supplemental Figure 7B).

In summary, our data reveals a heterogeneous mixture of cells in the *AML1-ETO* patients. Patient s220 showed more heterogeneity between Dx and Re compared with *AML1-ETO* s914. Interestingly in both patients, we found cells with a signficantly elevated LSC17-score that are predominantly in the G1-phase. These cells appear to be at the origin of other cell populations that develop/branch in a way that is sample and stage specific. The signature genes for LSCs might be potentially therapeutic targets to improve the efficiency of AML treatment.

## Discussion

To gain insight into the heterogeneity between AML subtypes and within Dx-Re pairs, we profiled the exome, gene fusions and single cell transcriptome of four *FLT3-*ITD and two *AML1-ETO* Dx-Re sample pairs. To our knowledge, this is one of the first studies analyzing Dx-Re pairs at an unprecedented depth of analysis. Clustering and differential expression analysis of single cell transcriptomes showed extensive intra- and inter-blasts heterogeneity. Genes that are differentially expressed between Dx and Re were highly patient-specific. Therefore, we chose a pairwise comparison and showed that differential expression is poorly predicted by altered somatic mutations in AML-associated genes. For example, one patient showed a pathway switch from AP-1 dependency at Dx to mTOR signaling at Re that appeared to be independent of altered somatic mutations, suggesting that clonal rearrangements are not causing the relapse^53^. In contrast, significantly altered mutations (e.g. loss of *NPM1* and *KIT*) in other patients were accompanied by minor transcriptional differences.

These results raise the question how the transcriptome of AML patients can be so drastically altered from Dx to Re in the absence of altered genomic aberrations? One possibility is that somatic mutations are gained or lost in regulatory regions that are not captured by exome sequencing. Alternatively, somatic mutations in genes that are currently not associated with AML may (collectively) contribute to therapy resistance. For example, in *FLT3-*ITD patient s3432 the clear separation of Dx and Re cells could be caused by de novo mutations in *FAT3, ITGB7, UBA2* and *SLC4A3*. Furthermore, the presence of quiescent LSC’s that escape conventional therapeutic interventions could explain recurrence in the absence of clonal rearrangements^14,54,55^. In agreement with this hypothesis, we detected transcriptionally similar LSC-like cells in the Dx and Re samples of the two otherwise distinct *AML1-ETO* samples. While the expression of these LSC populations is similar at Dx and Re, their differentiation trajectories are remarkably different.

Our study is based on few Dx-Re pairs, but nevertheless reports important findings that strongly indicate differences in underlying resistance mechanisms that are not exclusively caused by clonal rearrangements. Leveraging rapid advances in single cell technology, future studies analyzing more cases at the current unprecendented depth can address whether LSCs are indeed clonally identical at Dx and Re and to what extent therapeutic interventions and epigenetic mechanisms drive these marked differences in gene expression. Such in depths knowledge obtained experimentally and bioinformatically will open novel avenues to prevent AML relapse.

## Acknowledgements

This study was supported by the Princess Maxima Center for Pediatric Oncology, Utrecht the Netherlands, grants from ZonMw/ Bundesministerium fur Bildung und Forschung (German) (BMBF; DRAMA 01KT1603); VALERE: Vanvitelli per la Ricerca; Campania Regional Government Technology Platform “Lotta alle Patologie Oncologiche”: iCURE; Campania Regional Government FASE2: IDEAL; MIUR, Proof of Concept POC01_00043; Campania Regional Government: POR Campania FSE 2014-2020 ASSE III. Y.Z is a PhD student in co-tutele from the Traslational medicine PhD program at Vanvitelli University. W.M and N.D.G are supported by the Italian National Operational Programme on Research 2014-2020 (PON AIM 1859703-2). This work was carried out on the Dutch national e-infrastructure with the support of SURF Cooperative. Thank the lab members for fruitful discussions and suggestions.

## Author contributions

Y.Z performed experiments; A.D and L.B provided WES data; Y.Z, P.S, W.M and H.S analyzed and interpreted data; A.D, P.B, N.A, N.D.G, K.D, H.D. S.M, J.M, L.A and L.B helped with data interpretation; Y.Z and H.S designed the research; Y.Z, P.S, W.M and H.S wrote the manuscript.

## Data Availability

The high-throughput datasets have been deposited in the European Genome-phenome Archive. The accession numbers for single cell RNA-seq, bulk RNA-Seq and Whole exome sequencing datasets are EGAD00001008373, EGAD00001008374 and EGAD00001008375, respectively.

