## Supplemental material, fig1-6, table1 and 3 for "Longitudinal single-cell transcriptomics reveals distinct patterns of recurrence in acute myeloid leukemia"

Yanan Zhai *et al*.

### Supplemental Methods

#### AML samples

In total, 6 paired Dx-Re bone marrow aspirates from adult AML patients, aged between 31 and 69 and diagnosed with *AML1-ETO* (n=2 low risk cases) or *FLT3*-ITD (n=1 intermediate and n=3 high risk) were analyzed from the AMLSG BiO Registry study (NTC 01252485). The phenotypic characterization (*AML1-ETO* and *FLT3*-ITD) was performed at the reference laboratory of the German and Austrian AML Study Group (AMLSG). All patients received intensive standard induction chemotherapy with cytarabine and an anthracycline (7+3 regimen) followed by high-dose cytarabine consolidation cycles. Patient characteristics are summarized in **Supplementary Table S1**. Informed consent for both treatment and biobanking of leukemia samples according to the Declaration of Helsinki was given by all patients. Approval was obtained from the ethical review board of the University of Ulm (Ethikkommission der Universität Ulm).

#### Cell preparation

Bone marrow aspirates from AML patients were processed using density gradient centrifugation to isolate mononuclear cells, viably frozen in medium containing 10% DMSO and shipped with dry ice. Cryopreserved AML samples were thawed at 37℃. After washing with cold PBS, cells were resuspended in 500µl PBS with 0.2% human serum and incubated for 10 minutes on ice. After centrifugation and removing the supernatant, cells were resuspended in 200µl cold PBS with 20µl PE-conjugated CD33 and 20µl APC-conjugated CD34 (eBioscience). Cells were stained for 30 minutes on ice, washed with cold PBS with 0.5% BSA and resuspended in 500µl PBS with 0.5% BSA and 1µl 7-AAD (eBioscience). After incubation for 10 minutes, CD33/CD34+ cells were sorted into 384-well plate (BioRad) containing well-specific primers (100nl, 0.75pmol/µl) and 5µl mineral oil (Sigma-Aldrich) on the BD FACS Aria cell sorter. After sorting plates were sealed and span down for 2 minutes at 2000g, snap frozen on dry ice and stored at -80℃ until use.

#### Single cell SORT-seq

We applied SORT-seq[1], a method that integrates single cell FACS sorting (Fluorescence-Activated Cell Sorter) with the CEL-Seq2 protocol[2] to measure gene expression at single cell resolution. External RNA Controls Consortium (ERCC) transcripts were spiked-in to detect empty wells, low quality mRNA and failed reactions. The frozen plates were centrifuged at 1400 RPM for 2 minutes at 4°C before processing. Processing of single cell plates included first strand synthesis and barcoding in the 384 well plate. mRNA from the single cells were pooled and batch amplified by in vitro transcription (IVT) (Invitrogen # AM1334). Sequence libraries were prepared with Phusion High-Fidelity Polymerase (NEB).

#### Bulk RNA-seq

Total RNA was extracted using Quick-RNA Microprep Kit (Zymo Research) according to the manufacturer protocol with DNaseI treatment. The RNA concentration was quantified with Qubit Fluorometer (Invitrogen). RNA libraries were prepared with KAPA RNA HyperPrep Kit with RiboErase (HMR) kit (Roche) following the manufacturer's recommendations. RNA-seq libraries were paired-end sequenced on an Illumina Nextseq 500 at an average depth of ~30M reads.

#### Detection of *FLT3*-ITD at diagnosis and relapse

The presence of *FLT3*-ITD was detected by DNA-based PCR followed by capillary electrophoresis. The detailed procedures were described previously[3].

### Data analysis

#### Sequencing and mapping

Single cell libraries were pair-end sequenced on an Illumina NextSeq500 at an average depth of ~30M reads per library and demultiplexed using bcl2fastq version 2.15.0.4 with default settings. We used STAR version 2.7.2b[4] to map the 42nt long read1 to human reference genome hg38. Next, we used UMI-tools[5] to reconstruct the gene by cell UMI count matrix from the BAM file.

#### Normalization, dimensionality reduction and cluster analysis

We used the Seurat v3[6] R-package for downstream analysis. First, low quality cells (genes detected < 500 or UMI count > 12,000 or mitochondrial UMIs > 30% or ERCC reads > 20%) were discarded (**Supplementary Fig. 1A)**. Ribosomal and mitochondrial genes were also discarded prior to normalization. Cells from all libraries were concatenated and $\log_2$ normalized. Next, we applied principal component analysis (PCA) on the 2,000 most variable genes to reduce the dimensionality of the dataset and retained the 50 components for cluster analysis and the identification of marker genes. Cluster analysis was run using the Louvain algorithm. Cluster markers were identified using the Seurat function *FindAllMarkers* with parameters *min.pct*=0.25, *logfc.threshold*=0.5 and *only.pos*=FALSE. Marker genes discriminating two clusters or Dx from Re cells were obtained usng the Seurat function *FindMarkers* with parameters *min.pct*=0.25, *logfc.threshold*=0.5, *min.diff.pct*=0.2 and *only.pos*=FALSE. All marker genes with adjusted p-value > 0.01 were discarded.

#### Whole exome sequencing analysis

We used the GATK toolkit version v4.2.0[7] to detect short somatic variants following the GATK best practices workflows "*Data pre-processing for variant discovery*" and "*Somatic short variant discovery (SNVs + Indels)"* with small modifications. Briefly, paired-end reads were aligned using BWA version 2.2.1[8], discarding reads with MAPQ < 20. PCR duplicates were marked using Sambamba 0.8.0 *markdup*[9] and base quality scores were recalibrated using the GATK functions *BaseRecalibrator* and *ApplyBQSR* . Variants were called using the *panel of normals* (PON) and *gnomAD* VCF file provided in the GATK resource bundle in two modes: Dx and Re as tumor samples and Cr as a germline control, or all three samples as tumor only. The rationale for this approach is that variants present (at low frequency) in the Cr sample (due to minimal residual disease) are sometimes discarded as germline. Variants were filtered with *FilterMutectCalls* and annotated with the Ensembl Variant Effect Predictor (VEP) version 104[10].

Variants were discarded when ANY of the following conditions were satisfied:

- The variant FILTER status was unequal to "PASS" or "slippage."
- The variant had less than 5 reads on the alternative allele (AD < 5) in the Dx, Re and Cr samples;
- The variant allele frequency was below 0.05 (VAF < 0.05) in the Dx, Re and Cr samples;
- The variant had a gnomAD allele frequency $\geqslant$ 1.0 x $10^{-3}$;
- The variant allele frequency did not change significantly between Dx vs CR or Re vs CR (p-adjusted $\geqslant$ 0.01; Fisher's exact test)
- The variant allele frequency at Cr exceeded 0.2 ($VAF_{CR} > 0.2$)

Mutations were visualized using the *maftools* R-package[12] and listed in supplemental table 2.

### Supplemental Figure Legends

**Supplemental Figure 1. Quality Control of single cell data.** (A) Violin plots depicting the detected number of genes (top) and unique transcripts (bottom) per cell. (B) Venn diagram showing the number of DEGs between the Dx and Re sample, for a pairwise comparison in the four *FLT3*-ITD patients. Very few DEGs are shared between patients. (C) Same as (B), for the two *AML1-ETO* patients.

**Supplemental Figure 2. Single cell landscape of Single cell landscape of *FLT3*-ITD patient s232.** (A) UMAP of Dx and Re cells for *FLT3*-ITD patient s232 colored by timepoint (top) or cell cluster (bottom). (B) Heatmap displaying the top 20 cluster marker genes. Color represents row normalized expression values. (C) Overrepresented GO terms (category: biological pathway) in cluster 1 (Dx) and 3 (Re). P-values: hypergeometric test (BH-corrected). (D) Gene expression of selected mTORC1 pathway members.

**Supplemental Figure 3. Single cell landscape of *FLT3*-ITD patient s292.** (A) UMAP of Dx and Re cells for *FLT3*-ITD patient s292 colored by timepoint (top) or cell cluster (bottom). (B) Heatmap displaying the top 20 cluster marker genes. Color represents row normalized expression values. (C) overrepresented GO terms (category: biological pathway) per cluster.

**Supplemental Figure 4. Single cell landscape of *FLT3*-ITD patient s2275.** (A) UMAP of Dx and Re cells for *FLT3*-ITD patient s2275 colored by timepoint (top) or cell cluster (bottom). (B) Heatmap displaying the top 20 cluster marker genes. Color represents row normalized expression values. (C) overrepresented GO terms (category: biological pathway) for clusters 1 (Dx) and 5 (Re).

**Supplemental Figure 5. Relapse cells of *FLT3*-ITD patient s3432 are associated with exocytosis.** (A) Gene expression of selected RAS-pathway members. (B) overrepresented GO terms (category: biological pathway) for clusters 4. (C) Selected genes associated with exocytosis. Color depicts relative expression; size depicts the relative number of cells for a which at least one transcript was detected. ** $p < 0.01$, *** $p < 0.001$, Wilcox rank sum / Mann-Whitney U test.

**Supplemental Figure 6.** Cells in cluster 6 (blue circle, s914 *AML1-ETO*) simultaneously express hematopoietic stem-/progenitor- (*CD34*), monocyte (*LYZ*), B-cell (*MS4A1*), erythrocyte (*HBB*) and cell cycle (*TOP2A, MKI67*) marker genes. This indicates that these cells are doublets or contaminated by ambient RNA and were discarded from further analysis. Color bar represents the expression level of corresponding genes.

**Supplemental Figure 7. Single cell landscape of *AML1-ETO* patient s220.** (A) Heatmap displaying the top 20 cluster marker genes. Color represents row normalized expression values. Marker genes shared between cluster 4 (Dx) and 5 (Re) are highlighted inside a black rectangle. (B) overrepresented GO terms (category: biological pathway) at Dx and Re.

### Supplemental Table Legends

**Supplemental Table1.** Clinical information and sequencing details of the patients

**Supplemental Table2.** Dynamic changes of mutations between Dx and Re (WES) and detected fusion genes from bulk RNA-Seq. Please see the individual Excel table.

**Supplemental Table3.** Characterization of *FLT3*-ITD at diagnosis and relapse.

**Supplemental Figure 1.**

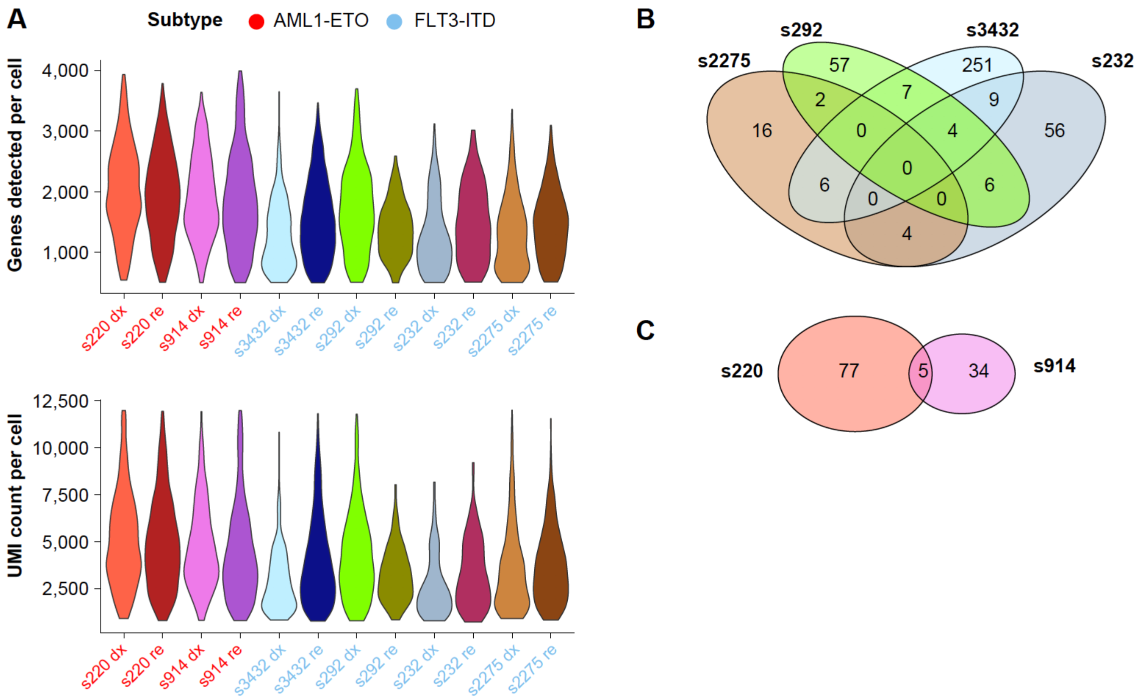

**Supplemental Figure 2 (s232).**

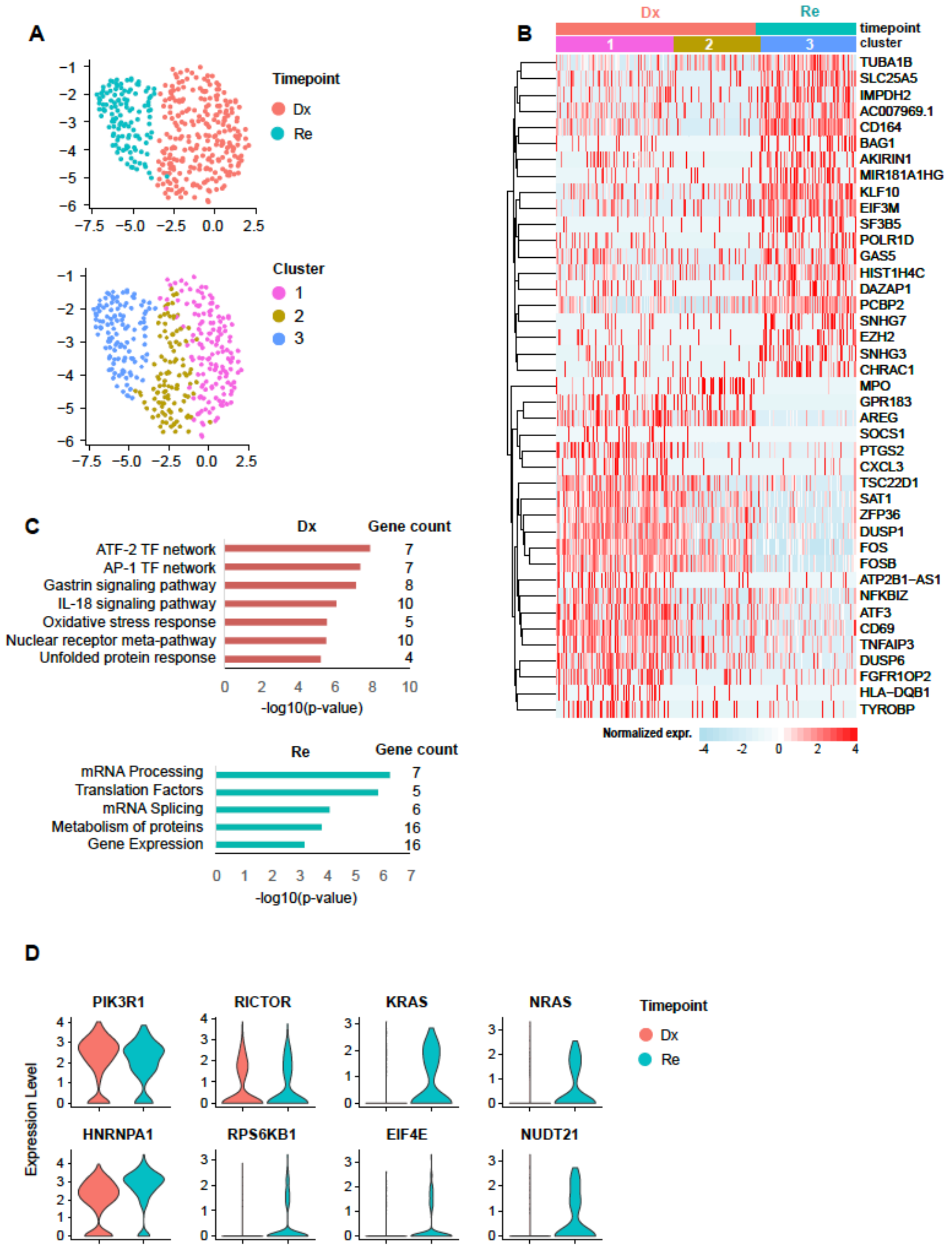

**Supplemental Figure 3 (s292).**

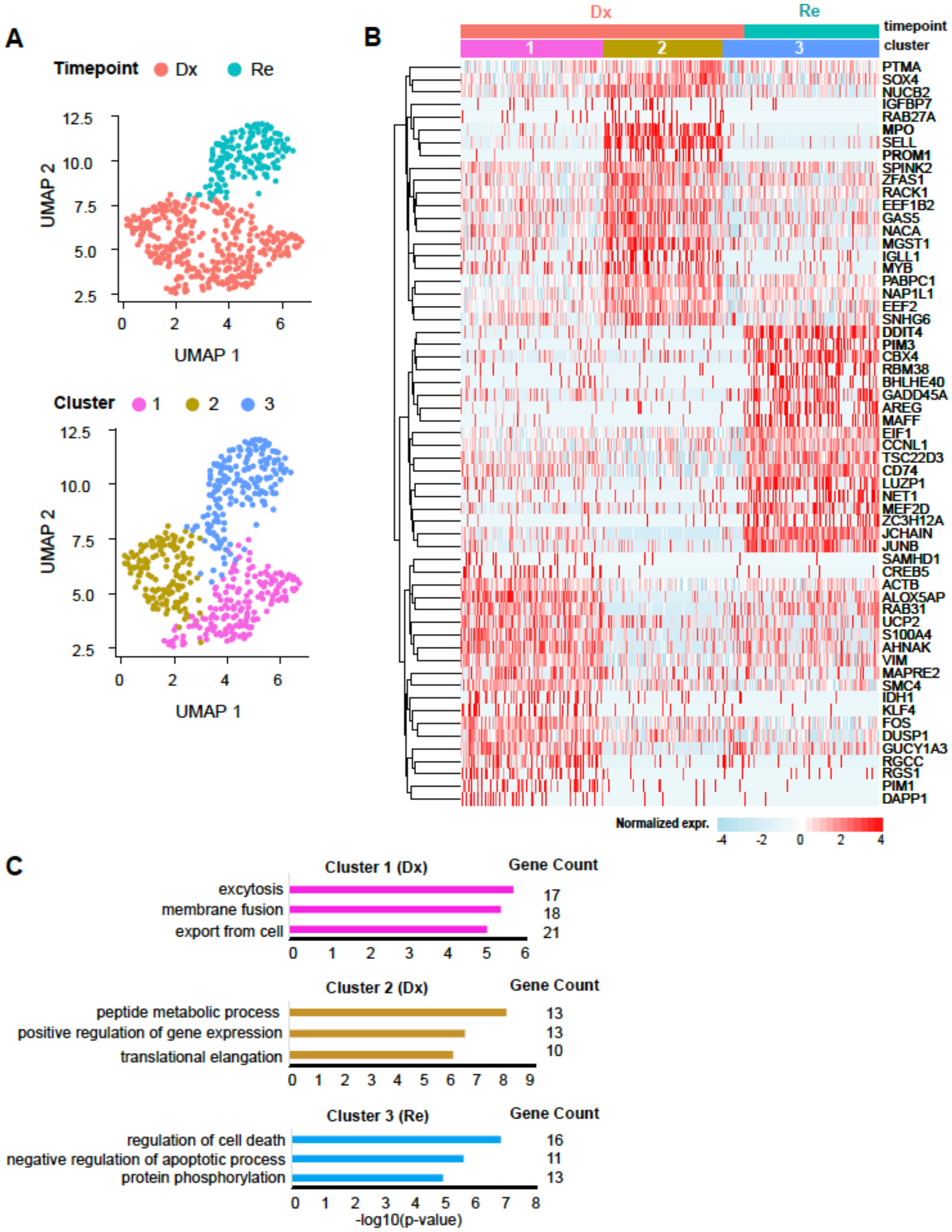

**Supplemental Figure 4 (s2275).**

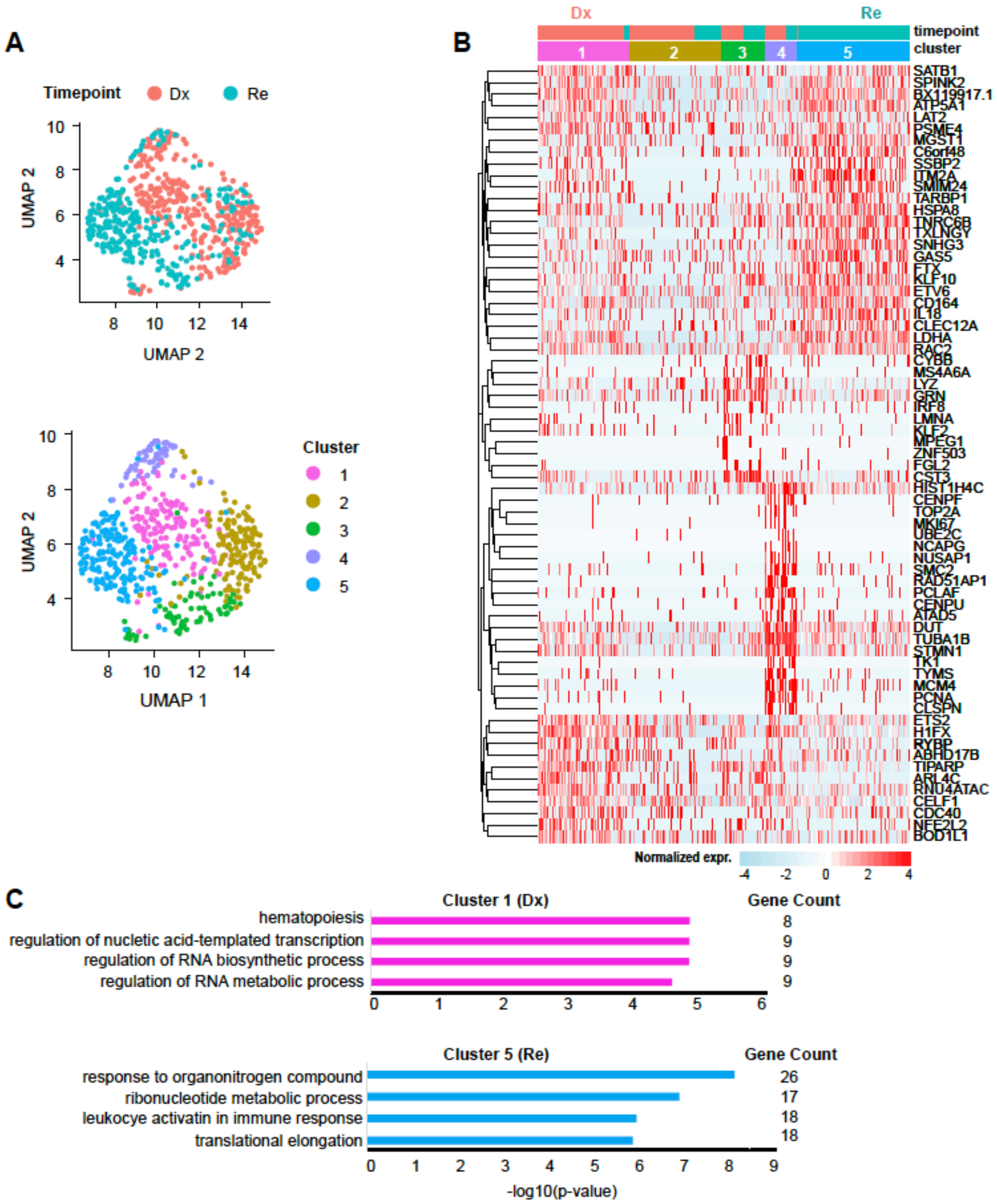

**Supplemental Figure 5 (s3432).**

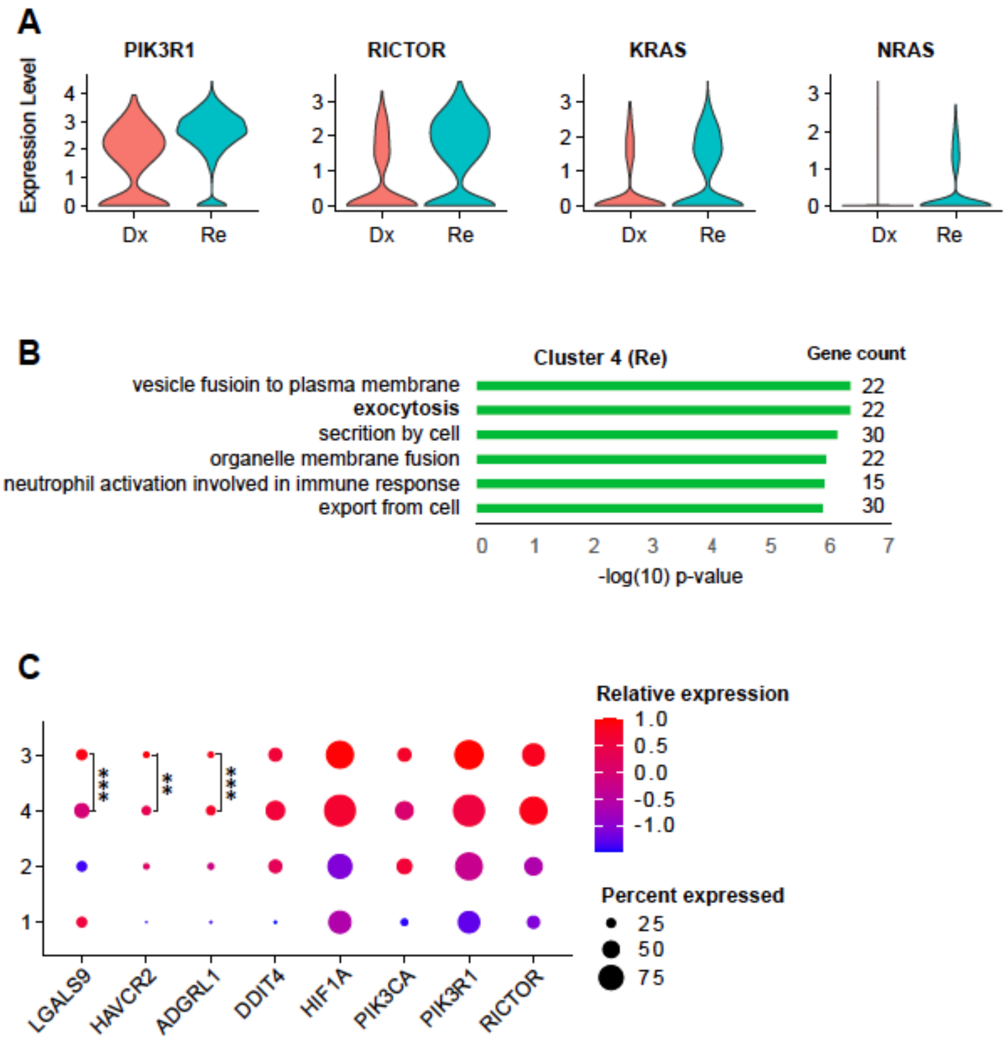

**Supplemental Figure 6 (s914).**

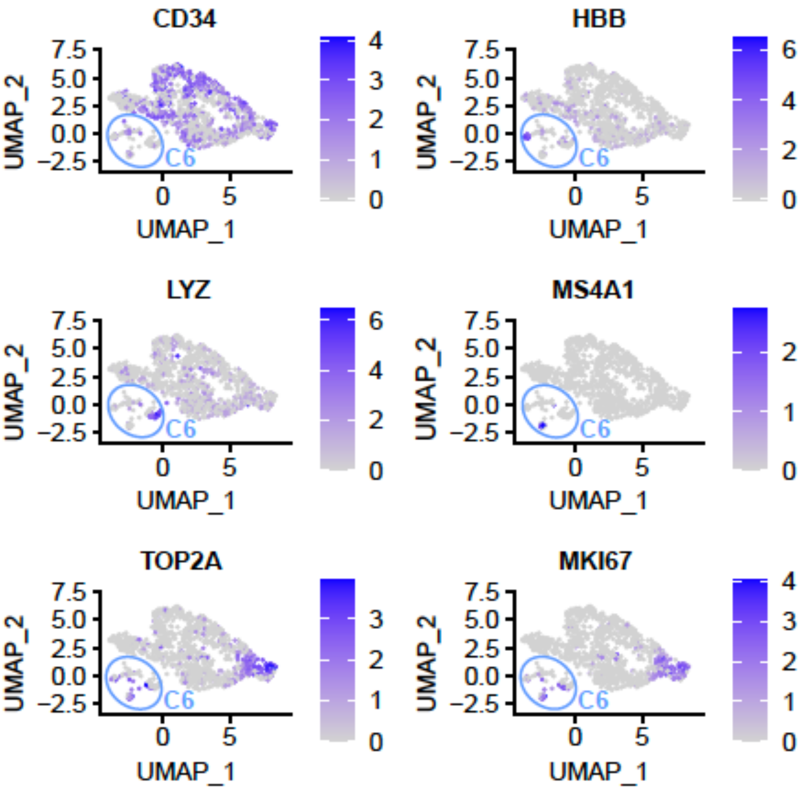

**Supplemental Figure 7 (s220).**

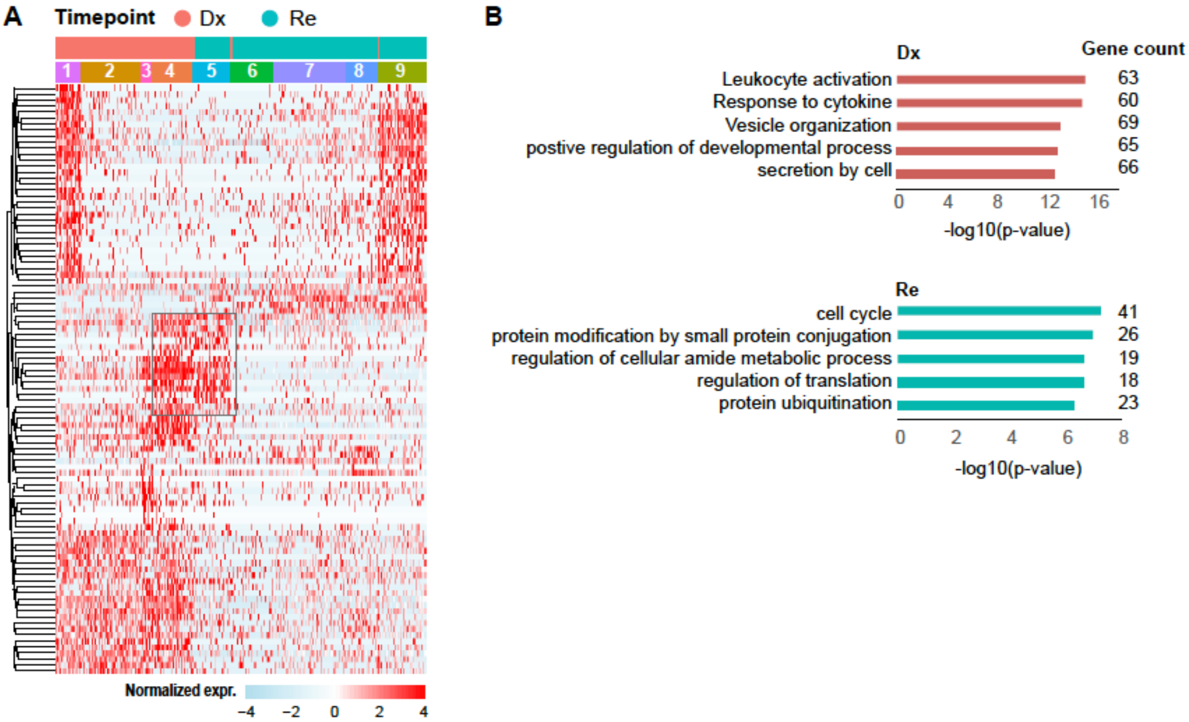

### Supplemental Tables

**Supplemental Table1.** Clinical information and sequencing details of the patients

| Patient ID | Gender | Age | Sample Source | | Blast% | | Enrichment | |
|---|---|---|---|---|---|---|---|---|
| | | | Dx | Re | Dx | Re | Dx | Re |
| **s220 *AET1-ETO*** | Male | 57 | BM | BM | 90% | NA | CD34 | CD34 |
| **s914 *AML1-ETO*** | Male | 31 | BM | BM | 30% | NA | CD34 | CD34 |
| **s232 *FLT3*-ITD** | Male | 53 | BM | BM | 80% | 90% | CD34 | CD34 |
| **s292 *FLT3*-ITD** | Male | 50 | BM | BM | 90% | 60% | CD34 | CD34 |
| **s2275 *FLT3*-ITD** | Male | 33 | BM | BM | 95% | 95% | CD34 | CD34 |
| **s3432 *FLT3*-ITD** | Female | 66 | BM | BM | 95% | 91% | CD33 | CD34 |

| Patient ID | Cell number after QC | | Counts per cell (Mean) | | Features per cell (Mean) | |
|---|---|---|---|---|---|---|
| | DX | Re | DX | Re | DX | Re |
| **s220 *AML1-ETO*** | 576 | 929 | 5 112 | 4 856 | 2 042 | 1 939 |
| **s914 *AML1-ETO*** | 314 | 688 | 4 713 | 4 411 | 1 951 | 1 851 |
| **s232 *FLT3*-ITD** | 249 | 123 | 2 653 | 3 289 | 1 278 | 1 497 |
| **s292 *FLT3*-ITD** | 324 | 152 | 4 224 | 3 261 | 1 726 | 1 380 |
| **s2275 *FLT3*-ITD** | 309 | 281 | 3 530 | 3 681 | 1 328 | 1 457 |
| **s3432 *FLT3*-ITD** | 509 | 1 158 | 2 895 | 3 956 | 1 299 | 1 563 |

BM: Bone Marrow; NA: not available

**Supplemental Table3.** Characterization of *FLT3*-ITD at diagnosis and relapse.

| Patient ID | Time point | FLT3-ITD allelic ratio | ITD same Dx/Rel 1=yes, 0=no | ITD loss (=FLT3-ITD negative at rel) 1=yes, 0=no | ITD change at rel 1=yes, 0=no | switch (insertion site, length) 1=yes, 0=no | loss of min 1 clone at rel 1=yes, 0=no | gain of min 1 clone at rel 1=yes, 0=no |
|---|---|---|---|---|---|---|---|---|
| **s232** | Dx | 0,398 | | | | | | |
| | Re | 0,744 | 0 | 0 | 1 | 0 | 1 | 0 |
| **s292** | Dx | 0,659 | | | | | | |
| | Re | 0,71 | 0 | 0 | 1 | 1 | 0 | 0 |
| **s2275** | Dx | 0,988 | | | | | | |
| | Re | 26,312 | 0 | 0 | 1 | 1 | 0 | 0 |
| **s3432** | Dx | 0,617 | | | | | | |
| | Re | 0,325 | 1 | 0 | 0 | 0 | 0 | 0 |
